# Evidence for Neolithic acquisition of the high pathogenic island by *Escherichia coli* followed by recent selection

**DOI:** 10.1101/2025.09.17.676721

**Authors:** Julie Marin, Erick Denamur, Olivier Tenaillon, François Blanquart

## Abstract

Many genetic elements involved in virulence and pathogenicity have been identified in commensal bacteria that are also opportunistic pathogens, but how these elements evolve is less well known. Understanding the evolutionary history of virulence elements requires genomes sampled in commensalism in healthy hosts, as genomes from bacterial infections are biased towards higher pathogenicity. Here, using a rare collection of human commensal *Escherichia coli* sampled in France from 1980 to 2020, we inferred the evolutionary history of the High Pathogenicity Island (HPI), a key virulence factor of *E. coli* with an uncertain evolutionary origin. The most likely scenario implied a horizontal transfer of the HPI from *Yersinia pestis*/*pseudotuberculosis* to the Enterobacterales, including *E. coli*, ∼4,700 years ago, coinciding with a period of increased population densities and human–animal interactions. The element subsequently spread in *E. coli*, with signs of positive epistasis with phylogroups B2 and F, and was positively selected in the last 100 years. Our work sheds light on the history and mode of evolution of virulence in an important opportunistic pathogen. It suggests the Neolithic period may have favored not only the emergence of zoonotic pathogens but also gene transfer between bacterial species, enriching the gene repertoire of major pathogens with potentially important implications on public health nowadays.

**Significance statement:** “Pathogenicity islands” are groups of genes that can be gained and lost on the chromosome of bacterial species and make them more pathogenic and virulent, but their evolutionary origin is unknown. We studied the origin and evolution of the High Pathogenicity Island of the major opportunistic pathogen *Escherichia coli* using molecular dating and phylogenetic analyses on very large genomic databases. We found that this island likely transferred from *Yersinia pestis*/*pseudotuberculosis* to *E. coli* 4,000 years ago, a period of increased population densities and intensified human–animal interactions, and was positively selected in the last 100 years. Thus, the origin of virulence can trace back to exchanges of mobile genetic elements that may be favored by particular circumstances in human history and have important consequences on human health nowadays.

## 1. Introduction

If human culture has left its mark on our genome—through the selection of genotypes aligned with cultural practices such as milk consumption—its impact on the microbes that colonize us has been even more pronounced. The advent of animal domestication increased human exposure to zoonotic pathogens, while the demographic consequences of agriculture transformed humans into a sustainable ecological niche for many infectious agents. More recently, the global spread of SARS-CoV-2 has illustrated how global patterns of hyper-connectivity shape pathogen transmission (du Plessis et al. 2021). Because pathogens have, in turn, shaped human history, they have long attracted intense scientific scrutiny. Theories of virulence (the severity of disease) and pathogenicity (the likelihood of causing infection) have become core topics in evolutionary ecology (Cressler et al. 2016). Yet one group of microbes has received far less attention: the silent majority of human-associated bacteria—commensals. How these organisms have responded to shifts in human culture remains poorly understood.

Filling this gap is important as many commensals are also opportunistic pathogens responsible for a substantial share of bacterial infections (Russo and Johnson 2003; O’Brien et al. 2009). While antibiotic consumption is known to select for resistance in these species (Austin et al. 1999; Baquero et al. 2002; Bell et al. 2014), much less is known about how cultural transitions have influenced their pathogenic potential and virulence. These traits are however genetically encoded, often clustered within pathogenicity islands—segments of DNA that carry virulence-associated genes (Hacker and Kaper 2000; Johnson 1991) and therefore subject to evolutionary variations. Accordingly, in *Escherichia coli*, one of the most consequential opportunistic pathogens, pathogenicity is highly heritable, with estimates of genetic contribution reaching 70% (Burgaya et al. 2023).

The evolutionary dynamics of *E. coli* virulence traits may be shaped by mutation, horizontal gene transfer and selection over the 100 Ma of evolution of this species (Ochman and Wilson 1987; Doolittle et al. 1996; Battistuzzi et al. 2004). The parallel acquisition of genetic elements associated to either intestinal or extra-intestinal virulence onto specific lineages may explain the formation of clones with greatly varying levels of intestinal and extra-intestinal pathogenicity through evolutionary history (Reid et al. 2000; Royer et al. 2023). Pathogenicity and virulence of *E. coli* measurably increased in France from 1980 to 2010 (Marin et al. 2022; Burgaya et al. 2023), but for the most part, how these elements evolved over the course of human history has not been much studied.

To investigate the evolutionary history of a common pathogenicity island, we focused on the High Pathogenicity Island (HPI), a genomic island coding for a siderophore-mediated iron capture system (yersiniabactin) and strongly associated with *E. coli* extraintestinal infections (Schubert et al. 2000; Schubert et al. 2004; Burgaya et al. 2023) and their severity (Galardini et al. 2020) but not with intestinal infections (Mey et al. 2021). In spite of the potentially important public health impact of this element, relatively little is known of its evolutionary history. This genomic island was first detected in *Yersinia pestis*, the causative agent of the plague (Carniel et al. 1992) and later found widely in the Enterobacterales family, including *E. coli*, *Klebsiella*, *Citrobacter* and *Salmonella* (Schubert et al. 2000; Carniel 2001). The HPI is flanked by the hybrid attachment sites, *attL* and *attR*, formed when the HPI integrates into the bacterial *attB* site (the *asn* tRNA gene) through site-specific recombination (Rakin et al. 2001). For most of the *E. coli* strains carrying the HPI, the 3ʹ-border including the *attR* site has been lost (Schubert et al. 1999). The origin of the *E. coli* HPI is unclear (Carniel 1999). The *E. coli* HPI is vertically and horizontally transmitted through conjugative transfer and homologous DNA recombination (Schubert et al. 2009), and increased in frequency in France from 1980 to 2000, suggesting selection (Marin et al. 2022). However, the dated evolutionary history of the HPI, the origin of the *E. coli* HPI, the mode of evolution and a quantification of selection on this element are all lacking.

Yet, the high degree of sequence identity of the HPI among the Enterobacterales and within *E. coli* clades suggests recent acquisitions of this island, opening the possibility of dating these events thanks to the wide availability of sequence data (Schubert et al. 1998). As large bacterial populations and close proximity might favor the transmission of genetic elements between bacterial species, transfers between *Y. pestis* and *E. coli* or other human commensals could be more likely during plague pandemic episodes. These episodes occurred throughout recent human history: *Y. pestis* is prevalent in ancient remains from the Neolithic and underwent a large-scale radiation between 6,000 and 5,000 years ago (Rascovan et al. 2019; Seersholm et al. 2024), and large outbreaks then burst during the Plague of Justinian (541–542) and the Black Death (14th century). The possibility that virulence elements cross species borders and rapidly evolve in humans raises several questions on the evolutionary dynamics of pathogenicity. What is the evolutionary origin of virulence genes? When were they introduced in *E. coli*, and how many times? Do they spread by selection or by horizontal gene transfer? Here, we used a rare dataset of human commensal *E. coli* collected from the 1980s to 2020 (Marin et al. 2022; Burgaya et al. 2023) to answer these questions for the HPI in Enterobacterales and *E. coli*.

## 2. Results

### 2.1 Evolutionary history of the HPI in Enterobacterales

#### 2.1.1 Support for a transfer of the HPI from *Y. pestis* to *E. coli*

The phylogenetic history of the HPI in Enterobacterales supported a scenario where the *E. coli* HPI have most likely originated from *Y. pestis/pseudotuberculosis*. To investigate the evolutionary history of the HPI in the Enterobacterales, we complemented a published commensal *E. coli* data set (423 strains) (Marin et al. 2022; Burgaya et al. 2023) with a recently published *Y. pestis* data set (147 strains) (Rascovan et al. 2019), and 2210 bacterial genomes from the RefSeq database including 52,208,856 sequences (Methods). We found this island in 26 additional species of Enterobacterales (**table S1**). Interestingly, HPI was found on 36 plasmid sequences in *Klebsiella* and 93 plasmid sequences in *Salmonella enterica*. In *E. coli*, we found the HPI in 236 strains over 423 and in 109 *Y. pestis* strains over 147. We removed branches and associated descendant tips (1197 tips) for which recombinant regions were longer than 5% of HPI sequence (**table S1**) and excluded the remaining recombinant regions (5,871 pb).

To decipher the evolutionary history of the HPI and the origin of the transfer to *E. coli*, we inferred a rooted phylogenetic tree of the HPI. We inferred the position of the root by maximizing the correlation between sampling dates and root-to-tip distances (Didelot et al. 2018). The placement of the root implied a scenario where the HPI originated from *Y. pestis* and spread to the Enterobacterales, possibly through *Y. pseudotuberculosis* (bootstrap value of 58%) (**figure 1)**. Although *Y. pestis* is a sublineage of *Y. pseudotuberculosis* (Bercovier et al. 1980; Achtman et al. 1999), it remains defined as a distinct species given its high virulence and unique ecological niche (Savin et al. 2019). The phylogenetic proximity of the two “species” (*Y. pestis* and *Y. pseudotuberculosis*) might partly explain the lack of resolution. The molecular rate was estimated to 4.82×10^−7^ SNPs per site per year (root-to-tips correlation), consistent with phylogenomic analyses of *E. coli* (Holt et al. 2013; Ben Zakour et al. 2016) and *K. pneumoniae* (Duchêne et al. 2016). This rooting scenario was strongly informed by the oldest sequences of *Y. pestis* HPI. Consequently, we explored alternative rooting scenarios, even though these alternative scenarios were not supported by our rooting method. An origin of the HPI within *Y. pseudotuberculosis*, with a subsequent spread to *Y. pestis* and the Enterobacterales, is also a likely hypothesis given the tight phylogenetic relationships of this species with *Y. pestis*. This scenario led to similar molecular rate of evolution (4.65×10^−7^) and inferred divergence times (**figure S2**). However, the molecular rates of evolution and the inferred divergence times were inconsistent with the literature for the other tested scenarios (see below). Specifically, the most recent common ancestor (MRCA) of the *Y. pestis* HPI was estimated to 56,612 years ago [95%HPD: 8388;217062] (root within *E. coli*) and 84,750 years ago [95%HPD: 8347;303951] (root within *K. pneumoniae*), i.e. >50,000 years before the emergence of the species 5,700-6,000 years ago (Rasmussen et al. 2015; Spyrou et al. 2018; Rascovan et al. 2019; Andrades Valtueña et al. 2022) (**figure S3 and S4**).

**Figure 1.**
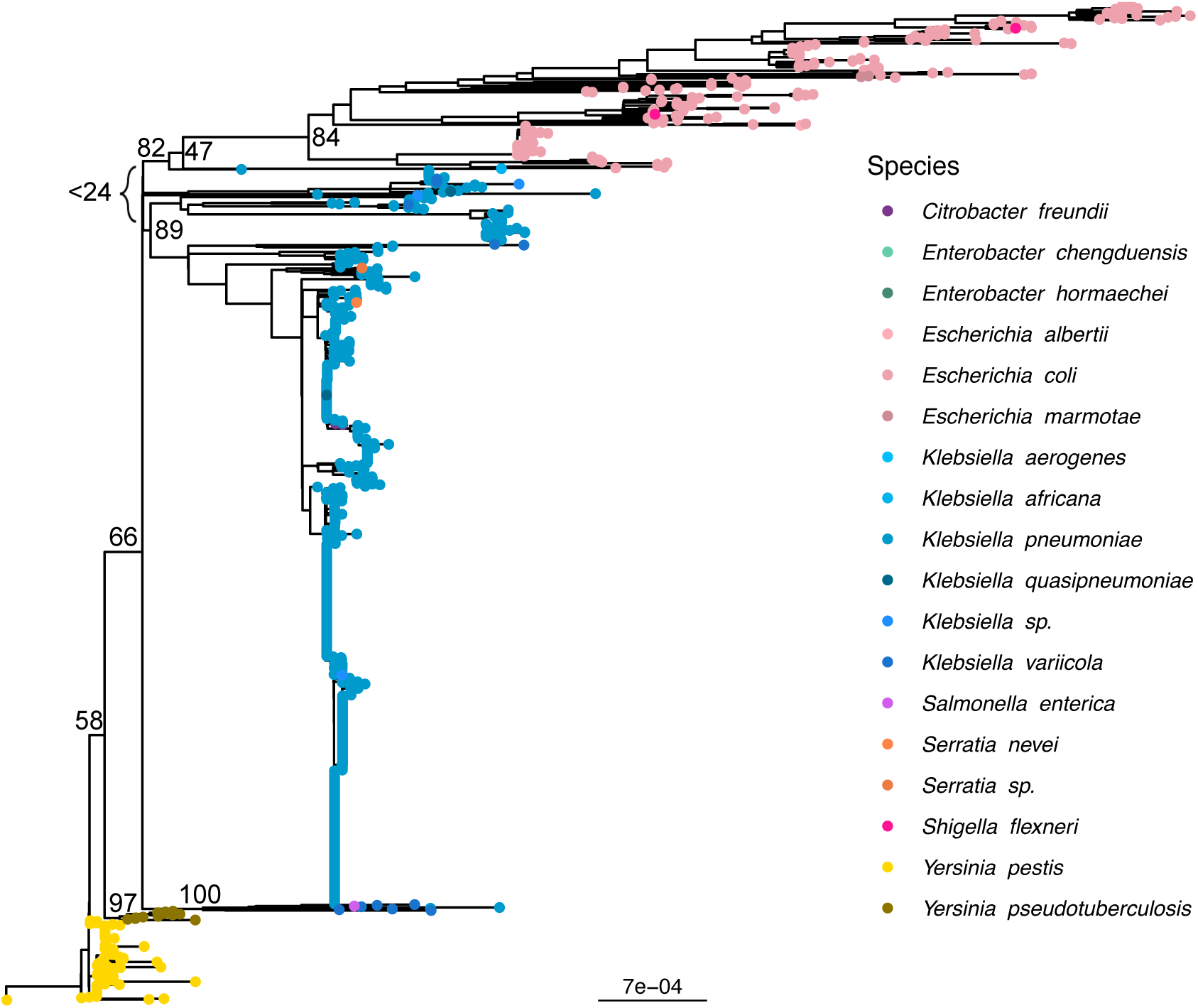
The HPI phylogenetic tree (IQ-TREE) rooted by maximizing the correlation between sampling dates and root-to-tip distances. The 1358 tips are colored by species. For clarity, we only show the associated bootstraps of the main clades.

The HPI in *E. coli* did not have a unique origin. However, most *E. coli* HPI sequences (221 *E. coli* and 2 *E. marmotae*/cryptic clade V) belonged to the same basal monophyletic group. Another *E. coli* HPI sequence was found in the large *K. pneumoniae* clade. Moreover, before removing branches and tips highly affected by recombination, 14 additional *E. coli* sequences were also interspersed within the *K. pneumoniae* clade, suggesting that the HPI can also be acquired by exchange with *K. pneumoniae* (**figure 1**).

The widespread distribution of the HPI among *E. coli* was mirrored in the diversity of coding regions directly surrounding the HPI (five coding regions upstream and downstream). More precisely, while flanking regions were highly conserved in the main HPI *E. coli* clade, the 15 strains scattered in the *K. pneumoniae* clade had several other surrounding regions.

Within the main *E. coli* clade, the DNA regions tRNA-Asn, *mtfA* and tRNA-Ser were found downstream of the integrase (*intA*) in 98.6% of the strains for which the HPI was not at the border of a contig (**figure 2 and table S2**). For the three remaining strains, two carried insertion sequences (ISSfl3) and one carried three hypothetical proteins. Upstream of the HPI, we found the same three hypothetical proteins for 99.6% of the strains for which the HPI was not at the border of a contig. The remaining strain carried an insertion sequence (IS1S).

**Figure 2.**
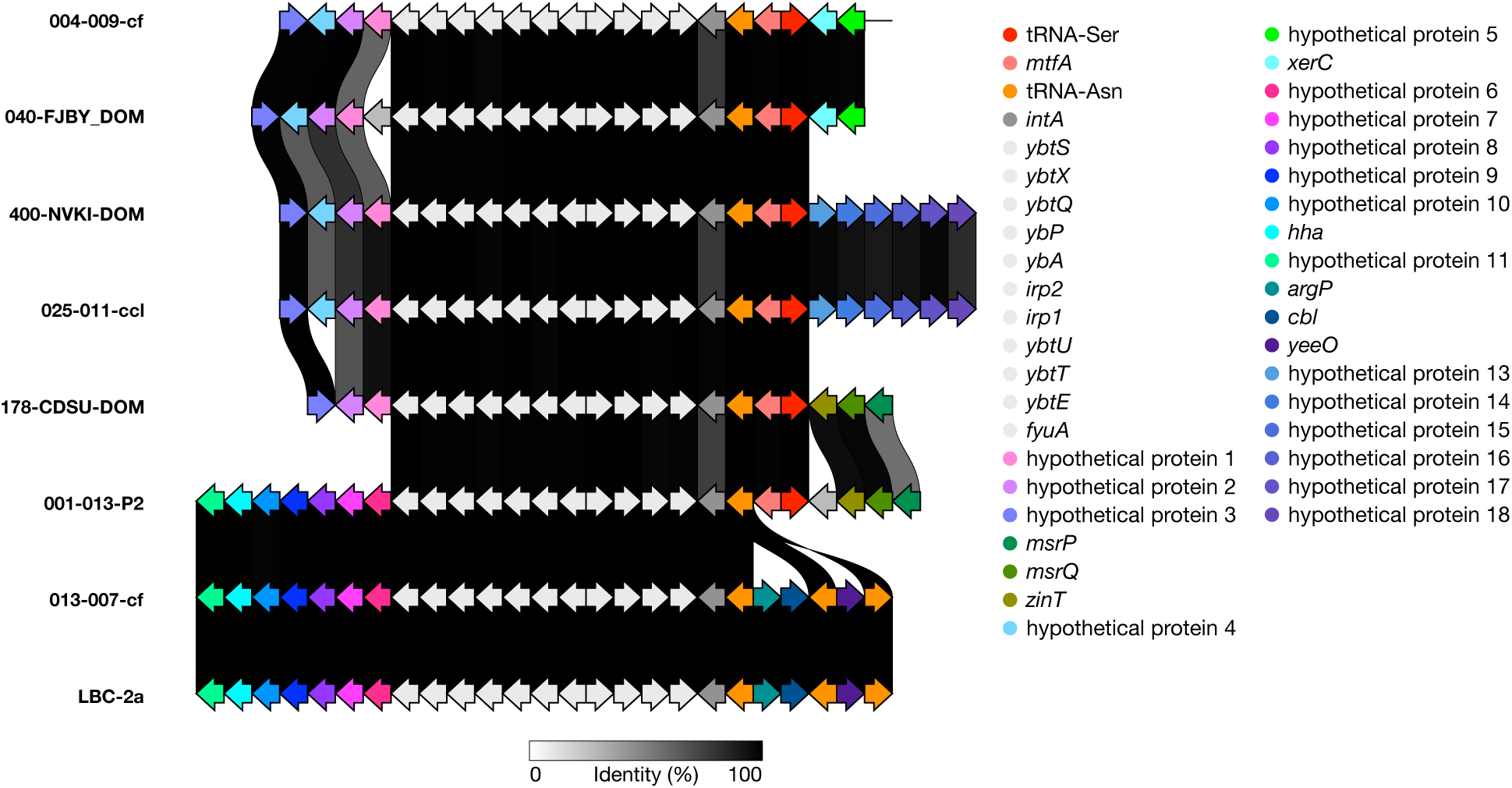
Gene synteny upstream and downstream of the HPI (in white). Genes are not shown to scale in order to visualise all of them. HPI of the last three strains, 001-013-P2 (kept after the removal of recombinant regions), 013-007-cf and LBC-2a are part of the HPI *K. pneumoniae* clade. We choose one to two representative sequences by main insertion sites (see **table S2**). All DNA regions are coding regions except two tRNAs (tRNA-Asn and tRNA-Ser). Homologous regions are shaded in grey.

Among the 15 strains found in the *K. pneumoniae* clade: 2 strains had the same coding regions (upstream and downstream) as the HPI of the *E. coli* clade (**table S2**); 7 strains had the same downstream coding regions as the HPI of the *E. coli* clade; 6 strains presented different coding regions (tRNA-Asn, *argP*, *cbl*). Upstream of the HPI, distinct coding regions from that of the HPI in the main *E. coli* clade were found for 80% of the strains (hypothetical proteins 6-11) (**figure 2**).

#### 2.1.2. Late Neolithic – Early Bronze age origin of the HPI in *E. coli*

We then dated the origin of the HPI in *E. coli* and *K. pneumoniae* using a subset of 389 sequences, and found the HPI was likely transferred to the Enterobacterales around 4700 years ago. To ensure convergence of molecular clock models in a reasonable computational time, we only kept five *K. pneumoniae* strains (randomly sampled); all strains with sampling date belonging to other species were kept in this analysis. The HPI of *E. coli* originated from the *Yersinia* clade including the oldest known strains, Gokhem2 (4900 years ago) and strains from Bronze Age (4800 - 3700 years ago) (Rascovan et al. 2019) (posterior probability of 1) (**figure 3**). We dated the origin of the HPI of the clade including all *E. coli* strains and *K. pneumoniae* strains to 4679 years ago [95%HPD: 3930;5351] (posterior probability of 0.80). The MRCA of the main *E. coli* HPI clade (221 sequences) was dated at 3978 years ago [3269;4766] (posterior probability of 0.80). The HPI in *K. pneumoniae* was slightly older, dated at 4186 years ago [2988;5245] (posterior probability of 0.58). Moreover, we found, based on the HPI sequences, that *Y. pestis* bronze age lineages and basal modern lineages diverged about 5212 years ago [4886; 5618], in line with a previous estimate of 5303 years ago based on whole genomes (Rascovan et al. 2019).

**Figure 3.**
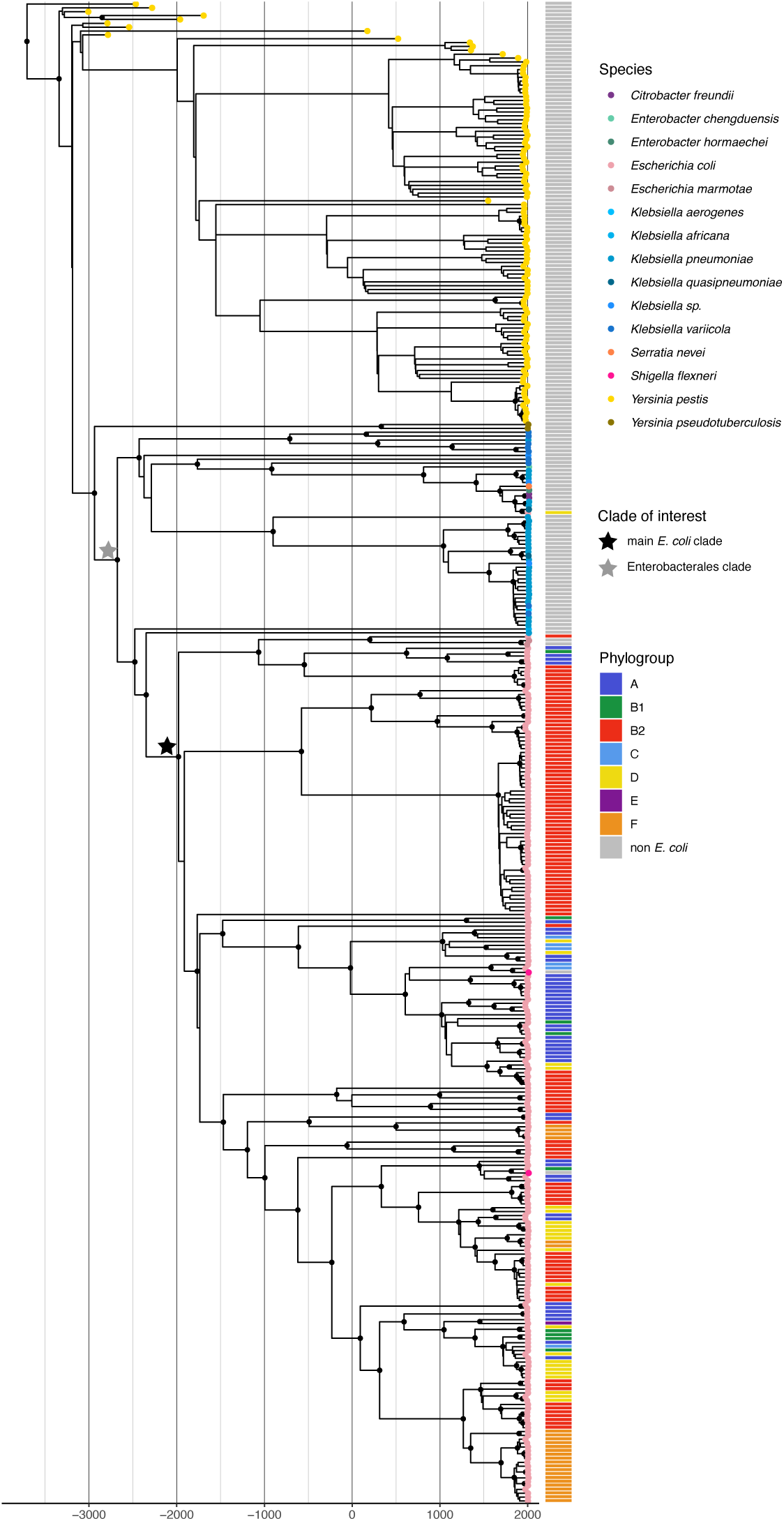
HPI dated tree (389 tips). Divergence times were inferred with BEAST2. Strains with no sampling date were removed and we only kept five strains from the *K. pneumoniae* clade. Black circles at nodes indicate posterior probabilities > 0.80. Stars correspond to two clades of interest, the main HPI *E. coli* clade (black) and the HPI *K. pneumoniae* clade (gray). HPD confidence intervals for node ages are shown in **figure S5**.

To further evaluate our estimates of divergence times we investigated the substitution rate for the three most abundant species in our dataset (*E. coli*, *K. pneumoniae* and *Y. pestis*). The molecular rate was estimated to 3.08×10^−7^ SNPs per site per year in *E. coli*, 1.59×10^−7^ in *K. pneumoniae* and 8.77×10^−9^ in *Y. pestis*. For the tree rooted with *Y. pestis*, our results were consistent with large-scale previous published work, where the molecular rate was inferred to 4.39×10^−7^ and 6.00×10^−7^ in *E. coli* (Holt et al. 2013; Ben Zakour et al. 2016), 2.99×10^−7^ in *K. pneumoniae* and *∼*10^−8^ in *Y. pestis* (Duchêne et al. 2016) (**figure S1A**). Our inferred rate for *E. coli* was also compatible to within-host rates estimated at 2.66×10^−7^ and 5.49×10^−7^ SNPs per site per year (Ghalayini et al. 2018; Condamine et al. 2025). However, rates of molecular evolution computed from the tree rooted in *K. pneumoniae* or *E. coli* were ∼100 fold slower in these two species and discordant with previous literature (**figure S1B and C**). These results further support our rooting within *Y. pseudotuberculosis/pestis*.

### 2.2 Evolutionary history of the HPI in *E. coli*

We next investigated the mode of evolution of the HPI within *E. coli* and found evidence for recent selection, and interaction between HPI and the genetic background. We reconstructed the frequency trajectory of the HPI over time and inferred the gain and loss rates to assess the strength of selection acting upon this virulence element. A pre-requisite to this analysis was to estimate the divergence times among *E. coli* strains in order to place the introduction of the HPI on this phylogeny and quantify selection at recent time scales.

#### 2.2.1 Dating the history of *E. coli*

One of the biggest challenges when dating bacteria trees is the small temporal span of calibration points (sampling times), as fossil data are lacking. To overcome this difficulty, we used *E. coli* strains sampled between 1930 and 1941 from reference collections, and added the information on the divergence time between *E. coli* and *Salmonella enterica*. We also used a dating method to accommodate the fact that rates of molecular evolution are much slower in the long-term (inter-species and inter-phylogroup divergences) than in the short-term (intra-ST divergences) (Ho et al. 2011; Duchêne et al. 2016). We adapted the time-dependent rates (“TDR”) method (Ho et al. 2005), using the tip dates of our 423 commensal *E. coli* strains sampled over 37 years, 24 *E. coli* strains from the Murray collection sampled between 1930 and 1941 (Baker et al. 2015), as well as the 102 Ma split between *S. enterica* and *E. coli* (Ochman and Wilson 1987; Doolittle et al. 1996; Battistuzzi et al. 2004) (**table S3**). We found a rapid decay of molecular rates as a function of clade age, from 1.78×10^−5^ in present time to 9.99×10^−8^ 1,000 years ago, comparable to the decrease in substitution rate observed for other human-associated bacterial species (Duchêne et al. 2016).

Using this method, we dated the *E. coli* species as well as cryptic clades IV and V to 34.71 Ma (95% confidence interval (CI) [33.42;35.40]) and *E. coli stricto sensu* (excluding cryptic clade I) to 9.72 Ma (95%CI [8.43;10.41]) (**figure S6 and table S3**). The four major phylogroups of our dataset started diversifying between 2.83 and 6.04 Ma, with phylogroup A dated at 3.18 Ma (95%CI [1.89;3.87]), phylogroup B1 at 2.85 Ma (95%CI [1.56;3.54]), phylogroup B2 at 2.83 Ma (95%CI [1.54;3.52]) and phylogroup D at 6.04 Ma (95%CI [4.75;6.73]). Regarding STs, we estimated the MRCA of ST131 to 1896.56 (95%CI [1815.15;1929.20], the MRCA of ST69 to 1967.68 (95%CI [1950.10;1978.06]), the MRCA of ST10 to 1599.25 (95%CI -479444.60;1814.32]) and the MRCA of ST95 to 1895.03 (95%CI [1811.20;1928.25]). Our time estimates were globally in line with previous estimates inferred from different dataset and methods (**table S3,** see Supplementary information).

#### 2.2.2 Evidence for selection on the HPI and epistasis with the genetic background

We then inferred the presence and absence of the HPI in the phylogeny of *E. coli*, and found a signal of positive selection for the HPI in the genomes (**figure 4A**). We used the dated phylogeny (TDR method) of the core genome (450 isolates) (**figure S6**). We then reconstructed 1000 random ancestral character histories (HPI presence/absence) using a Markov chain Monte Carlo approach and computed the HPI frequency through time, starting from a frequency of 0% at the time of acquisition by the main *E. coli* clade 3978 years ago (**figure 3**). We excluded the 15 strains for which the HPI was scattered through *K. pneumoniae* because they were probably governed by distinct, independent evolutionary dynamics. From this date on, lineages gained the HPI at a rate of 5.49×10^−2^ CI per year (CI [5.43×10^−2^; 5.54×10^−2^]) and lost it at 8.65×10^−2^ per year (CI [8.57×10^−2^;8.72×10^−2^]).

**Figure 4.**
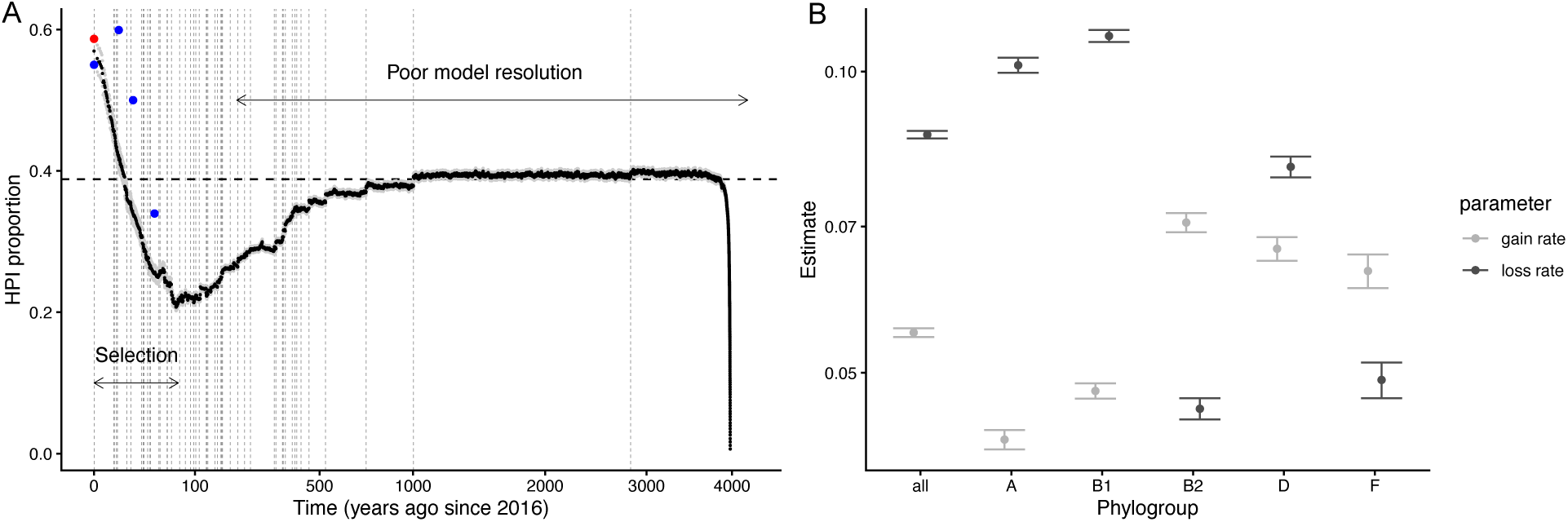
**(A)** HPI proportion through time (with square root transformation of the x-axis). The presence of HPI along branches, since its introduction, was computed from 1000 character histories sampled from the posterior probability distribution (95%CI in gray). The dashed vertical gray lines indicate the time of the first branching event for each ancestral lineage. The dashed horizontal black line shows the expected ratio at equilibrium (q01/(q01+q10)). HPI frequency in *E. coli* collections are illustrated by blues dots (CEREMI in 2016, Coliville in 2010, PAR/LBC/ROAR in 2001 and VDG in 1980). The HPI frequency at present time when considering all tips of the 54 clades together is indicated by the red dot. **(B)** HPI gain (q01) and loss (q10) rates estimates (with 95% CI) for *E. coli* tree and for the five main phylogroups (detailed results are presented **table S6**).

Overall, the HPI increased in frequency from 0% at the acquisition time to 59% in 2016. For most of the time span, corresponding to the stem branches connecting the crown of each ancestral lineage to the HPI introduction date, we had limited information. As a consequence, our model inferred a rapid spread to the expected equilibrium frequency reflecting the balance between acquisition and loss of the element, but not the HPI frequency over that period. As we entered more recent times and sampled within lineage diversification, we gained some resolution in estimating the HPI frequency through time. We found that the HPI increased in frequency over the last century due to faster branching rate of lineages carrying it, suggestion selection. Similar results were obtained with the lower and upper bound of the 95%HPD HPI acquisition date (**table S4**).

Here, the interpretation of positive selection on the HPI relies on the assumption that we sampled the whole diversity of the *E. coli* species. Alternatively, the increase in frequency of HPI could also be explained by sampling bias: if *E. coli* circulates in various host species, transfers across host species are rare and the HPI is less prevalent in other species than in human *E. coli*, then it is a possibility that clades with low frequency of HPI are not adapted to human. Apparent selection on the HPI could thus be explained by source-sink dynamics and not by direct selection on the HPI. However such source-sink dynamics would only produce observed patterns if animals were the primary hosts and humans the secondary hosts of HPI-free lineages, which is inconsistent with current epidemiological and ecological knowledge on *E. coli* (Tenaillon et al. 2010). To further test if the same selection trend was observed on a subset of strains strongly associated to human, we restricted our analysis to B2 strains (Touchon et al. 2020). We found a similar increase in frequency in recent times suggesting faster branching rate for lineages carrying the HPI and hence selection (**figure S7**).

The HPI was unevenly distributed among phylogroups (**table S5**). Its prevalence was 92.0% and 89.3% in the B2 and F phylogroups respectively, but only 38.4% and 18.3% in A and B1 respectively. We tested whether different trends existed between the five main phylogroups (A, B1, B2, D and F). Interestingly, for B2 and F strains the rate of loss was lower than the rate of gain (t-test, p-values < 2×10^−16^) (**figure 4B**). Moreover, the loss rate in these two phylogroup was also lower than for other phylogroups or for *E. coli* considered as a whole (pairwise t-test with Hochberg correction, p-values < 2×10^−16^). Both analyses suggested that the HPI loss and acquisition differed between phylogroups, more precisely that the HPI was more stable in B2 and F phylogroups (**figure 4B and table S6**).

## 3. Discussion

It was uncertain whether the *E. coli* HPI originated from *K. pneumoniae*, *Y. pestis*, or *Y. pseudotuberculosis* (Carniel 1999). Although future ancient DNA sequences may refine this inference, our analyses indicated that the HPI was probably transferred from *Y. pestis*/*pseudotuberculosis* to *E. coli* and other Enterobacterales approximately 4,700 years ago, coinciding with a period when plague was widespread in southern Scandinavia (Rascovan et al. 2019; Seersholm et al. 2024). Thus, the Neolithic is a period not only more favorable to the emergence of zoonotic diseases, but also possibly to cross-species gene transfer in bacteria. The HPI was then selected much more recently, with potential consequences on the incidence and severity of *E. coli* infections in modern times.

Thanks to a rare large dataset of *E. coli* genomes isolated from their natural habitat—the gut of healthy human hosts—we reconstructed the evolutionary history of this species, a pivotal result for the detection of recent selection acting on the high-pathogenicity island within the species. Our study contrasts with the frequent focus on strains from extraintestinal infections which exhibit lower genetic diversity (Kauffmann 1947; Denamur et al. 2021; Burgaya et al. 2023), and harbor more virulence genes (Clermont et al. 2017; Kallonen et al. 2017; de Lastours et al. 2020). To date the major events in the history of *E.* coli, we adapted the time-dependent rate method (TDR). This method enabled to model the systematic trend in molecular rate from very ancient times (the divergence between *S. enterica* and *E. coli* more than 100 Ma ago), to the past decades (*E. coli* strains sampled over the last 30 years), respectively reflecting the substitution and mutation rates (Ho et al. 2005). Globally, our dating is consistent with the literature, both at old (*E. coli sensus stricto*) and recent evolutionary timescales (STs) (**table S3** and Supplementary information).

The Neolithic was characterized by increased population densities and contacts with animals (Müller et al. 2016) which may have facilitated the emergence in humans of pathogens such as *Mycobacterium tuberculosis* (Hershkovitz et al. 2008), some strains of *S. enterica* (Key et al. 2020), and *Y. pestis* (Andrades Valtueña et al. 2022). We found that the HPI gene may have been introduced around 4,700 years ago in the Enterobacterales with support for a *Y. pestis*/*pseudotuberculosis* origin. Around 5,000 years ago, the plague was widespread in southern Scandinavia, with a prevalence of the order of 17% (Seersholm et al. 2024) This period also coincided with the large dispersal of Late Neolithic and Early Bronze Age lineages of *Y. pestis* across the European continent (between ∼4700–2800 BP), likely facilitated by increasing human mobility and trade networks (Andrades Valtueña et al. 2022; Swali et al. 2023). The origin of HPI in *Y. pestis* is supported by the available molecular data. An origin in *Y. pseudotuberculosis*, an enteropathogenic zoonotic bacterium, leads to similar dates of introduction in *E. coli* and *K. pneumoniae* and molecular rates of evolution also consistent with the literature (Figure S2) (Nakamura et al. 2016; Reinhardt et al. 2018). Such an origin is not supported by currently available data but is difficult to preclude entirely for lack of ancient *Y. pseudotuberculosis* sequences. These findings nevertheless suggest that the Neolithic transition, characterized by increased population densities and intensified human-animal interactions, may have provided favorable conditions to horizontal gene transfer among bacterial species.

The most common version of the HPI in *E. coli* diversified around 4000 years ago and spread in different phylogroups and clones of *E. coli* by conjugative transfer whereby the excised HPI is mobilized to a new recipient either trapped by a transmissive *asn* tDNA-carrying plasmid or autonomously as an ICE called ICE*Ec1* (Schubert et al. 2004). Vertical transfers are explained by the deletion of the 3ʹ-border that includes the hybrid attachment site *attR* in the vast majority of strains, hindering HPI excision (Schubert et al. 1999). The recent increase in frequency of the HPI could be explained by selection causing clonal expansion of the clades that carry this element. This may be caused by coincidental selection at the within-host level (Le Gall et al. 2007). For instance, dietary changes could favor strains with enhanced iron acquisition systems. Alternatively, as iron acquisition systems are associated with longer residence times (Östblom et al. 2011), between-host selection on longer residence times is possible. Such selection could be modulated by the frequency of disturbances of the microbiota, whereby reduced perturbations may favor longer-residence, more pathogenic genotypes (Morel-Journel et al. 2025).

Lastly, phylogroup-specific selective constraints shaping the HPI acquisition and loss may explain the preferential distribution of the HPI in B2 and F phylogroups. The HPI is rare in B1 (18%) and abundant in B2 and F (∼90%) in accordance with earlier results (Nowrouzian et al. 2009). Moreover, the loss rate of the HPI was smaller in phylogroups B2 and F than for other phylogroups. These results are in line with previous evidence for the role of epistatic interactions in the emergence of virulence in *E. coli*, with selection of lineage-dependent specific associations of virulence-associated genes (Royer et al. 2023). Alternatively, it is possible that phylogroups inhabit distinct ecological niches and that HPI is under stronger positive selection in the niches of B2 and F phylogroups (Morel-Journel et al. 2025).

Our work presents two limitations. Firstly, there remains an inherent, large uncertainty in dating when calibration points are absent (**figure S8**). This is apparent in the very wide confidence intervals for the dates of ancient STs (**table S3**). Very ancient *E. coli* sequences (dating 1000s of years) would help clarify the relationship between evolutionary rates and time, but as of now such sequences are not available. Our inference of recent evolutionary rates relies on the rate of evolution inferred in the STs with a molecular clock signal (**figure S9**) and are compatible with rates inferred for the close species *K. pneumoniae*, and with the relationship between rate and time across bacterial species (Duchêne et al. 2016). The relatively rapid recent rate of evolution in recent times (of the order of 10^−5^ substitutions per site per year) is driven by ST69. Relatedly, precisely dating the origin of the HPI was possible only thanks to the availability of ancient *Yersinia* sequences. Such ancient sequences were conserved in the dental pulp of individuals infected thousands of years ago, and are unfortunately not available as of now for opportunistic pathogens which rarely cause infection. It may be possible to obtain a correct order of magnitude of the age of transfer for other genes thanks to the molecular clock, provided the rate of evolution of these genes is not too variable. Dating the origin of accessory genes is critical for our analyses: in the absence of information on the date of the transfer, one could have erroneously inferred a much more ancient origin of HPI based on the pervasiveness of the gene in the *E. coli* phylogeny (Didelot et al. 2012; Royer et al. 2018). Secondly, our hypothesized root of the HPI phylogeny was essentially informed by the molecular diversity of sequences with respect to sampling dates. The rooting method selects the roots that best aligns the tips of the phylogeny with the sampling dates, favoring a root close to the most ancient sequences. In the absence of an outgroup, the root could also be inferred with molecular clocks (*e.g.* BEAST (Drummond et al. 2006)) which are robust to small deviations from the clock-like behavior (Mai et al. 2017; Tria et al. 2017). However, the large variation in substitution rate existing among, and also within, species of our dataset, prevented us from using such methods (Duchêne et al. 2016). Other algorithms, such as “midpoint rooting” (Farris 1972) or those based on non-reversible substitution models (Minh, Hahn, et al. 2020; Naser-Khdour et al. 2022), would favor roots that homogenize the branch lengths, irrespective of sampling dates. This placement of the root led however to unrealistic divergence times (see Material and Methods).

Investigating the evolutionary dynamics of pathogenicity in commensal bacteria that are also opportunistic pathogens is a promising research avenue. Several bacterial species with large burden—such as *Streptococcus pneumoniae*, *Staphylococcus aureus*, and *K. pneumoniae*—are primarily commensals. If pathogenicity is, at least in part, determined by bacterial genetic factors, then it is subject to evolutionary change, likely shaped by the ecology of these organisms in their commensal lifestyle. To better understand these dynamics, future studies should focus on bacterial genomes sampled from healthy hosts, systematically compare them with genomes derived from infections, and could detect selection acting on pathogenicity and virulence-associated elements. Several other virulence genes are located on pathogenicity islands—regions of the chromosome with distinct GC content and that may have originated from other species. This is the case of the siderophores aerobactin (*iuc* operon), Sit-encoded iron transport system (*sit* operon) and salmochelin (*iro* operon) (Royer et al. 2023); and the genotoxin colibactin (*pks* operon) (Chagneau et al. 2022). Applying these methods systematically to these accessory genes could unravel the mode of evolution of bacterial species and their pangenomes and the recent selective pressure acting upon them.

In conclusion, available evidence is consistent with a transfer of the HPI from *Y. pestis*/*pseudotuberculosis* to *E. coli* and other Enterobacterales in the late Neolithic. The HPI spread by selection over the past decades and may now cause increased pathogenicity with important consequences on disease burden in modern times. This calls for more systematic research on the mode of evolution of bacterial species and their pangenomes and its interplay with the evolutionary history of humans.

## 4. Methods

### 4.1. Evolutionary history of the HPI in Enterobacterales

The HPI is a 36 to 43 kb region encoding the siderophore yersiniabactin (Schubert et al. 2000) composed of 12 genes (*fyuA*, *ybtE*, *ybtT*, *ybtU*, *irp1*, *irp2*, *ybtA*, *ybtP*, *ybtQ*, *ybtX*, *ybtS* and *intA*). We conducted an exhaustive search of the HPI (extracted from the strain 040-FJBY_DOM and excluding the highly variable *intA*) across bacterial species (excluding *E. coli* and *Y. pestis*) in GenBank using Blast 2.14 (https://blast.ncbi.nlm.nih.gov/Blast.cgi) against the database RefSeq (accessed on the 2025/11/06, Release 232 including 52,208,856 sequences), with the following parameters: 90% identity and 90% coverage. Using the core nucleotide BLAST database, which contains non redundant sequences, we excluded identical entries. For *Y. pestis* strains, we used the whole-genome assemblies with sampling dates (gathered from Rascovan et al. 2019). There were two exceptions: for two ancient strains, BlackDeath and Gokhem2, we used draft genomes mapped to *Y. pestis* CO92 (Rascovan et al. 2019). Commensal *E. coli* strains were gathered from six previously published collections: VDG in 1980 (n=53) (2), ROAR in 2000 (n=50) (3), LBC in 2001 (n=27) (4), PAR in 2002 (n=27) (4), Coliville in 2010 (n=246) (5) and CEREMI in 2016 (n=20) (Burgaya et al. 2023). We identified gene positions with ABRicate (Seemann 2022), using 50% coverage and 90% identity except for *irp1* for which we used 0% coverage and 90% identity because it was often found on several contigs. We excluded strains with less than 10 HPI genes (**table S7**).

#### 4.1.1 Phylogenetic history of the HPI in Enterobacterales

##### Rooting the HPI tree

We aligned the HPI sequence (from *fyuA* to *ybtS*) with mafft v7.487 (Katoh and Standley 2013) and built the phylogeny (2555 strains) with IQ-TREE v2.4.0 (1000 bootstraps and GTR+F+I+G4 model selected via the *ModelFinder Plus* option) (Kalyaanamoorthy et al. 2017; Minh, Schmidt, et al. 2020). We used ClonalFrame to detect recombinant events (Didelot and Wilson 2015). We removed descending tips of branches for which more than 5% of the alignment was affected by recombination (1197 tips). Next, we excluded the remaining recombinant sites (5,871 sites, 19% of the alignment). We took advantage of the wide range of sampling date of our dataset (from - 3000 to 2023) to root the tree by maximizing the correlation between sampling dates and root-to-tip distances, using the R function ‘initRoot’ (R package BactDating (Didelot et al. 2018)). In doing so, we assumed that the position of the root is mainly informed by sampling dates rather than by branch lengths which largely depend on substitution rates which are highly variable in our dataset (Duchêne et al. 2016). To test how the favoured rooting depend on the dates of our samples and particularly the fact that the oldest strains are exclusively from *Y. pestis*, we tested a more balanced species temporal range. We restrained the data-set to strains sampled between the oldest *Y. pestis* strains (2006) and the younger non *Y. pestis* strains (1980), resulting in 129 strains. Here, when maximizing the correlation between sampling dates and root-to-tip distances, the root was placed within *E. coli*. This placement of the root would lead to substitution rates (**figure S1C**) and divergence times inconsistent with the literature (see results).

Alternatively, we used another rooting approach using non-reversible models (Naser-Khdour et al. 2022). We compared the log-likelihood of trees being rooted on every branch of the ML tree with IQ-TREE v2.1.4 (UNREST substitution model and--root-test option) (Minh, Schmidt, et al. 2020). However, we did not find strong support for any of the root placements tested. For instance, the most likely root placement was supported by a bootstrap proportion of 0.35 (RELL approximation) (Kishino et al. 1990). The low support for root placement was likely the consequence of the limited number of informative sites (929 sites) which could inform the placement of the root (Naser-Khdour et al. 2022). Moreover, this method slightly favored a root within the *K. pneumoniae* clade. This scenario would imply a transfer from *K. pneumoniae* to *E. coli*, and later to *Y. pestis*. Similarly, to the rooting within *E. coli* (see above), this alternative rooting lead to substitution rates (**figure S1C**) and divergence times not supported by previous studies (see results).

##### Identification of coding regions surrounding the HPI

We first extracted from the gff files (Prokka output (Seemann 2014)) the 5 DNA regions (CDS and tRNA) upstream and downstream the HPI (including the *intA* gene) of one random strain. We searched those 10 DNA regions around the HPI in all strains with ABRicate (Seemann 2022) with more than 90% coverage and 95% identity. These steps were repeated with a new strain until we identified the 5 coding regions upstream and downstream the HPI for all the 235 strains with HPI, except if the contig ended. Next, we visualize the synteny of the main insertion sites (shared by at least 5 strains) with clinker (Gilchrist and Chooi 2021). For eight selected strains (one or two by main insertion site), we examined 5000 bp before and after the HPI. We set the minimum alignment sequence identity to color a link between two genes at 50%.

#### 4.1.2 Dating the *E. coli* HPI

The estimation of divergence time of the HPI was performed with the program BEAST2 v2.7.3 (Bouckaert et al. 2019) with the following parameters: optimized relaxed clock, substitution model GTR+I+G, at least 200 million generations with sampling every 5000 generations and a burn-in of 10 percent. We constrained the monophyly of the two first diverging clades from the root to preserve the optimal rooting. The sample times were used to calibrate the tips. We checked the chain convergence (effective sample sizes (ESS) > 200). We resampled the trees every 150,000 steps with LogCombiner and summarized the resulting trees with TreeAnnotator. We only kept five randomly sampled *K. pneumoniae* strains to ensure reasonable computational time and removed strains with no sampling date, resulting in a 389-tip timetree. We ran two replicates with two tree priors, coalescent with constant population and coalescent with Bayesian skyline. The best-fitting model was determined by computing Bayes factor from the marginal likelihood estimates calculated by Nested Sampling method (1 particle) (Russel et al. 2019) (**table S8**). The best-fitting model (coalescent with Bayesian skyline) was used in the subsequent analyses.

We employed the same method to infer divergence times of the HPI tree rooted within *Y. pseudotuberculosis*. Finally, to explore the alternative (not supported) scenarios, we also dated the divergence times of HPI tree rooted within the *K. pneumoniae* clade (non-reversible model) and the HPI tree rooted within the *E. coli* clade (root-to-tip method with same age ranges among *Y. pestis* and other species) following the same method.

### 4.2 Evolutionary history of the HPI in *E. coli*

#### 4.2.1 Evolutionary history of *E. coli*

We dated the *E. coli* phylogenetic tree, including 423 *E. coli* from our commensal collections, 24 *E. coli* strains from the Murray collections to increase the temporal span and three *S. enterica* as the outgroup. We used the least-square dating (LSD) and the time-dependent rate (TDR) methods, using sampling dates and the divergence between *S. enterica* and *E. coli* to calibrate the tree (see Supplementary information for more details).

#### 4.2.1 Selection on the HPI in commensal *E. coli*

Because the HPI was acquired by *E. coli* after the emergence of the species, we cut the tree into 54 sub-trees corresponding to the largest trees for which the mrca (most recent common ancestor) was younger than our estimation of the HPI introduction date (4197 BP) in the main *E. coli* clade. We added a stem from each crown to the HPI introduction date by binding an outgroup branch to each subtree with a tip in the present. By default, the HPI was set to “absent” at the tip of this outgroup, except when all tips of the clade were in state “HPI absent” (25 clades), in which case we set the outgroup to “present” to be able to include all subclades in the analysis. We computed the transition rates (HPI gain or loss) by estimating the ancestral character states along the tree using a continuous-time Markov chain model (R package phytools (Revell 2024)). We first estimated the parameters q01 (gain rate) and q10 (loss rate) for each of the 54 subclades where the two states were present, covering 89% of the total number of tips. We estimated the most likely rate matrix Q for each subclade and computed the mean q01 (gain rate) and q10 (loss rate). Next, for each subclade, we sampled 1000 evolutionary histories of the character from their posterior probability distribution. We calibrated the prior distribution of each parameter (q01 and q10) as a gamma distribution with the means of q01 and q10 across subclades, and the variance set to 1×10^−03^. To reach convergence (ESS > 200), we ran the MCMC algorithm over 100,000 generations with sampling every 100 generations (and retained the simulations after a burn-in period of 1000 generations). To get estimates for the five main phylogroups (A, B1, B2, D and F) we extracted clade specific results depending on their phylogroup membership. Next, we computed and plotted the HPI frequency through time over the 1000 character histories of each subclade. Additionally, we computed the HPI frequency through time for B2 strains only by applying the same methodology to the B2 phylogroup which included 11 sub-clades.

## Supplementary information

### Estimation and analyses of divergence times of the *E. coli* tree

#### 1. Data-set

The collection of commensal *E. coli* strains was sampled over 37 years (1980-2016). In order to improve the temporal sampling, we used 24 sequences from the Murray collection with samples ranging from 1930 to 1941 (Baker et al. 2015) (**table S9**). From the 50 genomic sequences available in the Murray collection, we selected sequences with available sampling time and excluded multiple variants per strain (when strain name, date on tube and origin were identical for two samples). Using independent lines of evidence and data-sets, the split between *S. enterica* and *E. coli* has been estimated to have occurred between 120 and 160 Ma (Ochman and Wilson 1987) and 102 Ma [57;176] (Battistuzzi et al. 2004). We assumed that the divergence occurred at 102 Ma because the most recent study used a larger molecular data set. To use this node as a secondary calibration point in our temporal analysis, we added three *S. enterica* sequences (accession number: CP023468, CP028169, CP054715) from Enterobase (Zhou et al. 2020).

The resulting 450 assemblies (423 *E. coli* from our commensal collections, 24 strains from the Murray collections and three *S. enterica*) were annotated with Prokka 1.14.6 (Seemann 2014). We then performed pan-genome analysis from annotated assemblies with Panaroo v1.3.0 with strict clean mode and the removal of invalid genes (Tonkin-Hill et al. 2020). We generated a core genome alignment spanning the whole set of core genes as determined by Panaroo and we generated the corresponding phylogenetic tree with FastTree v2.1.10 (Price et al. 2010).

#### 2. Estimation of divergence times

We inferred the phylogenetic history of *E. coli* from the core alignment (3 224 199 nt) for 447 *E. coli* (ingroup) and 3 *S. enterica* sequences (outgroup). We generated a maximum likelihood phylogenetic tree with FastTree v2.1.10 (Price et al. 2010). We checked that there was a molecular clock signal in our data-set (*E. coli* tree *stricto sensu*) using tips calibration (r=0.18, p-value=9.75 × 10^−5^).

We estimated the divergence times of the species *E. coli* using two different methods, least-squares dating method (LSD) and time-dependent rate method (TDR), with *S. enterica* sequences as the outgroup. We did not date the *E. coli* tree with the software BEAST for two reasons. First and mainly, we needed to date the whole tree (450 tips) to evaluate the selection strength on the HPI in *E. coli* with trait-dependent diversification models. Even after removing non-informative sites, the large size of the alignment (717 587 nt) prevented us from reaching convergence in reasonable time. Second, the molecular clock models implemented in BEAST is thought to describe phylogenies spanning small evolutionary scales, such as viral phylogenies (Lartillot et al. 2016), because the rates on branches are drawn from a single underlying parametric distribution such as a log-normal (Lepage et al. 2007; Rannala and Yang 2007; Drummond et al. 2012). We thus resorted to alternative methods described below.

##### a. Least-squares dating method

The first method, LSD, is a normal approximation of the Langley-Fitch model (To et al. 2016). The number of substitutions along each branch of the tree is modelled by a Poisson distribution with a single rate (strict molecular clock) (Langley and Fitch 1974). The error variance can accommodate uncorrelated variation of rates across branches. We used tip calibrations and we set the root calibration at 102 Ma. We ran the program LSD2 with the following parameters: rooted tree, constrained mode (to ensure that the date of every node is equal or smaller than the dates of its descendants), the application of variances on branch lengths (to compensate the overconfidence given on very short branches), sequence length and 1000 bootstraps.

##### b. Time-dependent rate dating method

We adapted a second dating method with time-dependent rate (TDR), in an attempt to reconcile the long-term and the short-term rates of molecular evolution (Ho et al. 2005). In essence, this method assumes no heterotachy (variation in lineage-specific molecular rates) and transforms the phylogenetic distances to temporal distances, while accounting for a directional trend in the rate of molecular evolution. Indeed, a negative relationship between the estimated rate and the depth of calibration has been observed in several datasets including humans and viruses (Howell et al. 2003; Ho et al. 2005; Santos et al. 2005; Gibbs et al. 2010). Among other factors, such as sequencing or calibration error, the presence of purifying selection is the most likely explanation (Ho et al. 2005). On very short timescales (*e.g.,* between successive generations), estimated rates include nearly all the genetic differences, approaching the spontaneous mutation rate (except mutations too deleterious to allow the isolation of the strain). At longer timescales (*e.g.,* between species), the estimated rate approaches the substitution rate. Substitutions are the mutations that get fixed in each divergent lineage through the action of natural selection and drift. Substitution rates are therefore usually much lower than the mutation rates (Ho et al. 2011). As a consequence, to simultaneously estimate deep (between phylogroups) and recent (within STs) divergence times, we need to formulate a model of rate variation that includes a directional trend towards faster rates closer to the present. The TDR method answers this problem by modelling the relationship between calibration age and rate as a decreasing exponential function as suggested by Ho et al. (Ho et al. 2005).

The relationship between calibration age and rate was generated as follows. We estimated the divergence times of five frequent STs (more than 6 strains) with a molecular clock signal (*i.e.,* positive relationship between root-to-tip distance and time): ST69, ST93, ST95, ST141 and ST452. To test the robustness of the molecular clock signal, we permuted sampling dates 1000 times within each ST. The observed number of STs with a molecular clock signal (5 STs) was reached in only 3.2% of the permuted pseudo-datasets (**figure S9**).

For these 5 STs, and for the whole data-set, we estimated the molecular rate of evolution as the slope of the relationship between root-to-tip distances and sampling dates. For the whole data-set, we used an additional point to estimate the slope corresponding to published estimates, setting the root at 102 Ma (Battistuzzi et al. 2004). For STs, the age of calibration was defined as the date of the MRCA, that is the date when root-to-tips distance was extrapolated to be 0. We obtained a negative relationship between calibration age and rate (**figure S8**). We note that random variability in root-to-tip distances uncorrelated with time would artifactually generate this relationship, as a steep slope will be associated with a recent MRCA. Here, the robust clock signal (**figure S9**), and the variability in the diversity of STs (Burgaya et al. 2023) should limit the impact of this artefact. Next, following the method described in (Ho et al. 2005), we fitted a vertically translated exponential curve to the relationship between calibration age and rate:

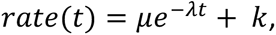

with *t* the calibration age, *μ* the instantaneous mutation rate (minus the rate of lethal mutations) and *λ* inversely proportional to the half-life of the rate decay. The constant term *k*, is a finite asymptote that vertically translates the curve and represents the evolutionary rate over long time periods. We fixed *k* to 1.51 × 10^−9^, our estimate of the molecular rate of evolution for the whole data set with the divergence between *E. coli* and *S. enterica* calibrated at 102 Ma. The confidence intervals of the parameters *μ* and *λ* were estimated with the R function ‘confint2’ (nlstools package (Baty et al. 2015)).

The TDR method is a rate varying method that is applied to a phylogenetic tree with equal root-to-tip distances for contemporary sequences (no heterotachy). To transform the phylogenetic tree so that contemporary sequences have the same root-to-tip distances, we used the LSD method with sampling dates only (To et al. 2016) and rescaled the tree to match the phylogenetic tree depth. Next, we estimated the molecular rates according to the TDR method as follows (Ho et al. 2005). The divergence date (in years) of two sequences (denoted *t_d_*) was estimated by solving the following equation for *t_d_*, given their sequence divergence (*d*):

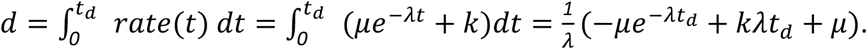

with the *μ*, *λ* parameters estimated from the analysis of the six STs described above and *k*, fixed to 1.51 × 10^−9^ as estimated from the whole dataset.

#### 3. Comparison of divergence times among methods

The TDR method accounts for the systematic temporal variability in the rate of evolution (Ho et al. 2005). Using this method, we dated the emergence of *E. coli stricto sensu* to around 9.72 Ma, consistent with a previous estimate of 9.0 Ma (95%CI [8.1Ma;9.9Ma]) using the rate of synonymous substitution (Reid et al. 2000). The TDR method also dated evolutionary events at intermediate timescales where previous existing methods cannot give reliable results: we dated the MRCA of phylogroup D at 6.04 Ma, A at 3.18 Ma, B1 at 2.85 Ma, and B2 at 2.83 Ma. Sequence types—largely monophyletic clones defined by seven gene alleles (Wirth et al. 2006)—generally span less than 100 years, with the notable exception of ST10 of phylogroup A which dates from 400 to 600 years ago. Our dating of ST was for the most part similar to previous studies (**table S3**). We dated the most recent common ancestor (MRCA) of ST131 to 1896.56 (95%CI [1815.15;1929.20], which is comparable to previous estimates, 1874 (95% highest posterior density (HPD) interval [1697;1951]) and around 1900 respectively (Kallonen et al. 2017; Arredondo-Alonso et al. 2025). We dated the MRCA of ST69 to 1967.68 (95%CI [1950.10;1978.06]), similar to previous estimates of 1956 (95%HPD [1935;1971]) and 1951 (95%HPD [1924;1969]) (Kallonen et al. 2017; Marin et al. 2022). A recent study including a more basal clade (unknown *fimH* allele) dated the ST69 MRCA around 1830 (Arredondo-Alonso et al. 2025). Our MRCA estimates of ST10 (1599.25 (95%CI [-479444.60;1814.32])) and ST95 (1895.03 (95%CI [1811.20;1928.25])) were younger than previous estimates which did not include the systematic increase in molecular rates toward the present (Kallonen et al. 2017; Marin et al. 2022; Arredondo-Alonso et al. 2025). The difference is particularly pronounced for the oldest ST (ST10), 222 years younger according to our estimate.

We also estimated divergence times with the “least-square dating” (LSD) method (**table S3**). The *E. coli* divergence date was similar to the TDR estimate, 40.53 Ma (95%CI [31.72;51.37]). Phylogroup MRCA were of the same order of magnitude but always older than TDR estimates. However, the ST MRCA estimates were much older with the LSD than the TDR method. For example, ST131 and ST69 were dated to 0.91Ma (95%CI [0.66Ma;1.22Ma]) and 0.46Ma (95%CI [0.34Ma;0.60Ma]) respectively. These discrepancies between the LSD and TDR methods are mostly explained by their rate estimation through time (**figure S10**). The model underlying LSD considers a constant rate of molecular evolution with a varying error term among branches to reflect the clock heterotachy (To et al. 2016), and cannot accommodate strongly and systematically time-varying rates contrary to the TDR method.

## Acknowledgments

We are very grateful to Nicolàs Rascovan for providing alignments and mapped sequences of *Yersinia pestis*. We thank Olivier Clermont for collection curation. We also thank the INRAE MIGALE bioinformatics facility (MIGALE, INRAE, 2020. Migale bioinformatics Facility, doi: 10.15454/1.5572390655343293E12) for providing computing resources. JM and FB were funded by the CNRS Momentum grant to FB.

## Data Availability Statements

No new data were generated or analyzed in support of this research. The code used to analyze the data is available here: https://github.com/j-marin/HPI.

