## Supplementary Figures for "Evidence for Neolithic acquisition of the high pathogenic island by *Escherichia coli* followed by recent selection"

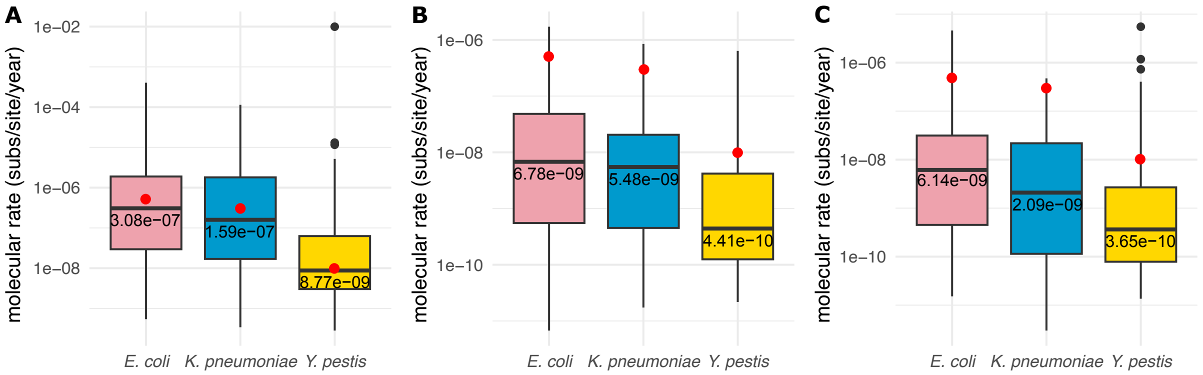


**Figure S1.** Molecular rate estimates (per branch, number of substitution/site/year) for the three main species, *E. coli* (in pink), *K. pneumoniae* (in blue) and *Y. pestis* (in yellow) computed from the comparison between the phylogenetic tree and the timed tree rooted in *Y. pestis* (A, main scenario), in *E. coli* (B, unsupported scenario) and in *K. pneumoniae* (C, unsupported scenario). Red dots indicate molecular rate estimates from the literature : $5.00 \times 10^{-7}$ for *E. coli* (Holt et al. 2013; Ben Zakour et al. 2016), $2.99 \times 10^{-7}$ for *K. pneumoniae* and $1 \times 10^{-8}$ for *Y. pestis* (Duchêne et al. 2016). Rates of evolution of the HPI are ~100-fold slower in *E. coli* and *K. pneumoniae*, and not consistent with the previous literature, when the root of the HPI tree is in *E. coli* (B) or *K. pneumoniae* (C).

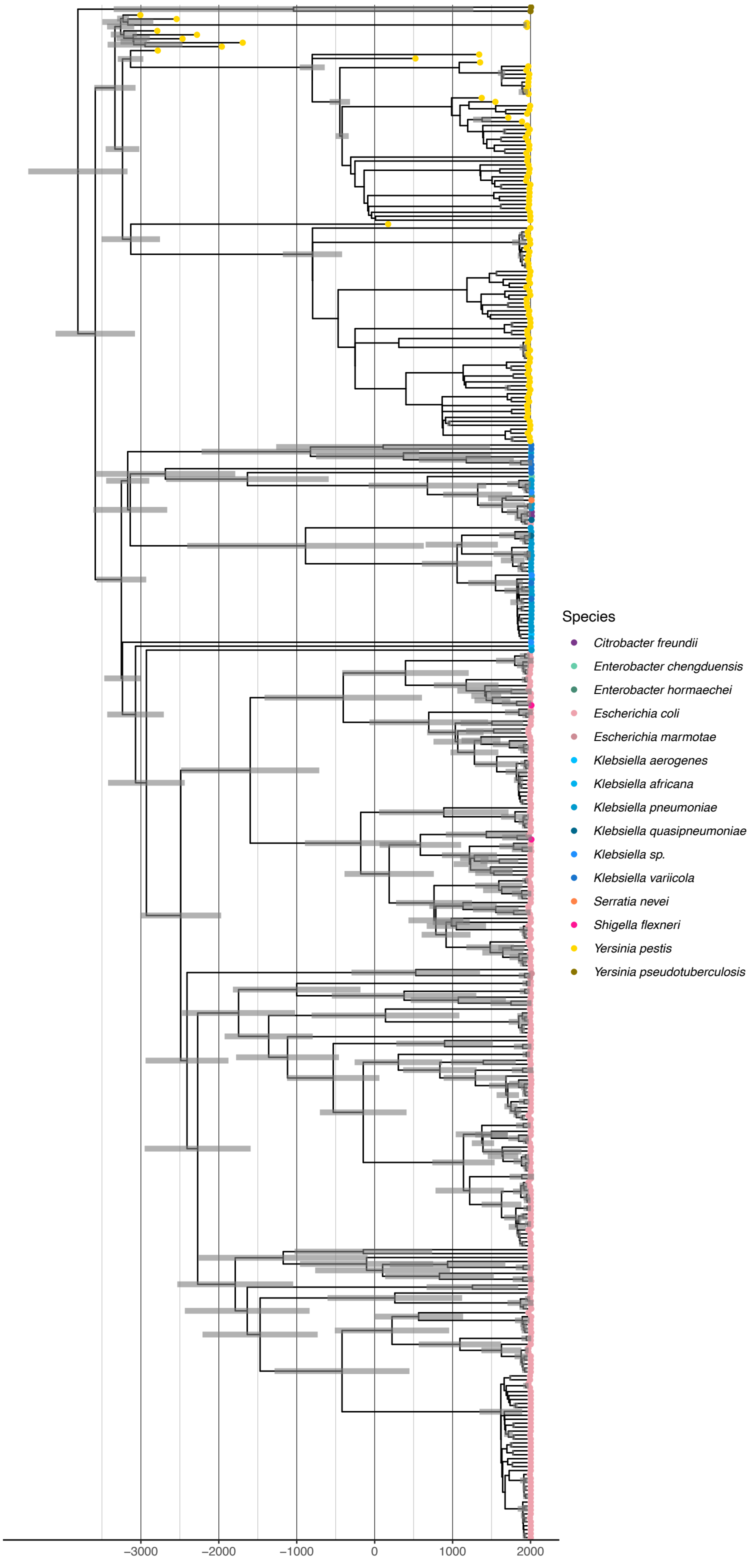


**Figure S2.** HPI dated tree rooted within *Y. pseudotuberculosis* (389 tips) built with BEAST2. HPD 95% confidence intervals for node ages are displayed as horizontal gray bars at each node.

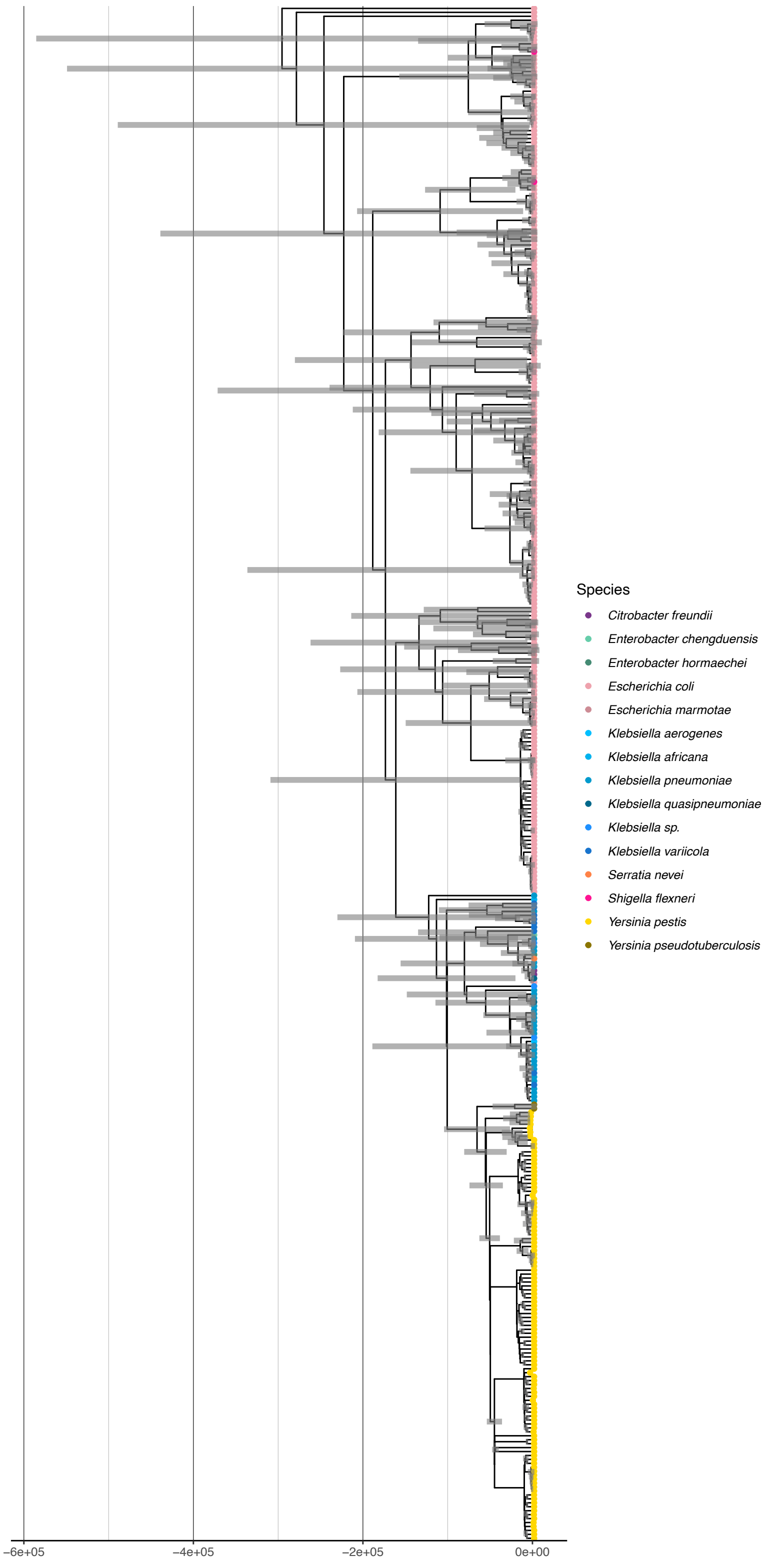


**Figure S3.** HPI dated tree rooted within *E. coli* (389 tips) built with BEAST2 (unsupported scenario). HPD 95% confidence intervals for node ages are displayed as horizontal gray bars at each node.

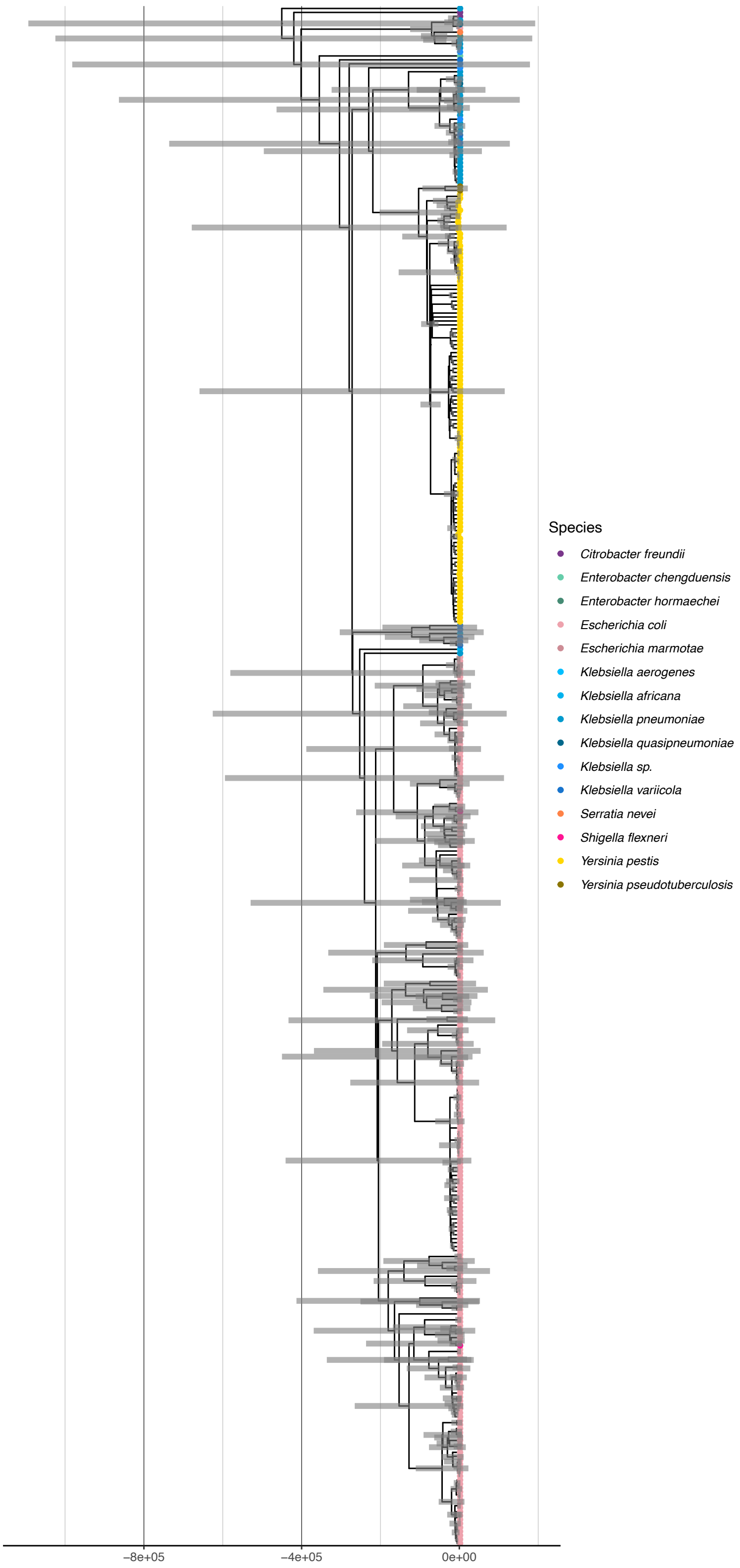
Species
Citrobacter freundii
Enterobacter chengduensis
Enterobacter hormaechei
Escherichia coli
Escherichia marmotae
Klebsiella aerogenes
Klebsiella africana
Klebsiella pneumoniae
Klebsiella quasipneumoniae
Klebsiella sp.
Klebsiella variicola
Serratia nevei
Shigella flexneri
Yersinia pestis
Yersinia pseudotuberculosis
−8e+05
−4e+05
0e+00

**Figure S4.** HPI dated tree rooted within *K. pneumoniae* (389 tips) built with BEAST2 (unsupported scenario). The scale of the HPD 95% confidence intervals for node ages is too small compared to branch lengths to be visualized.

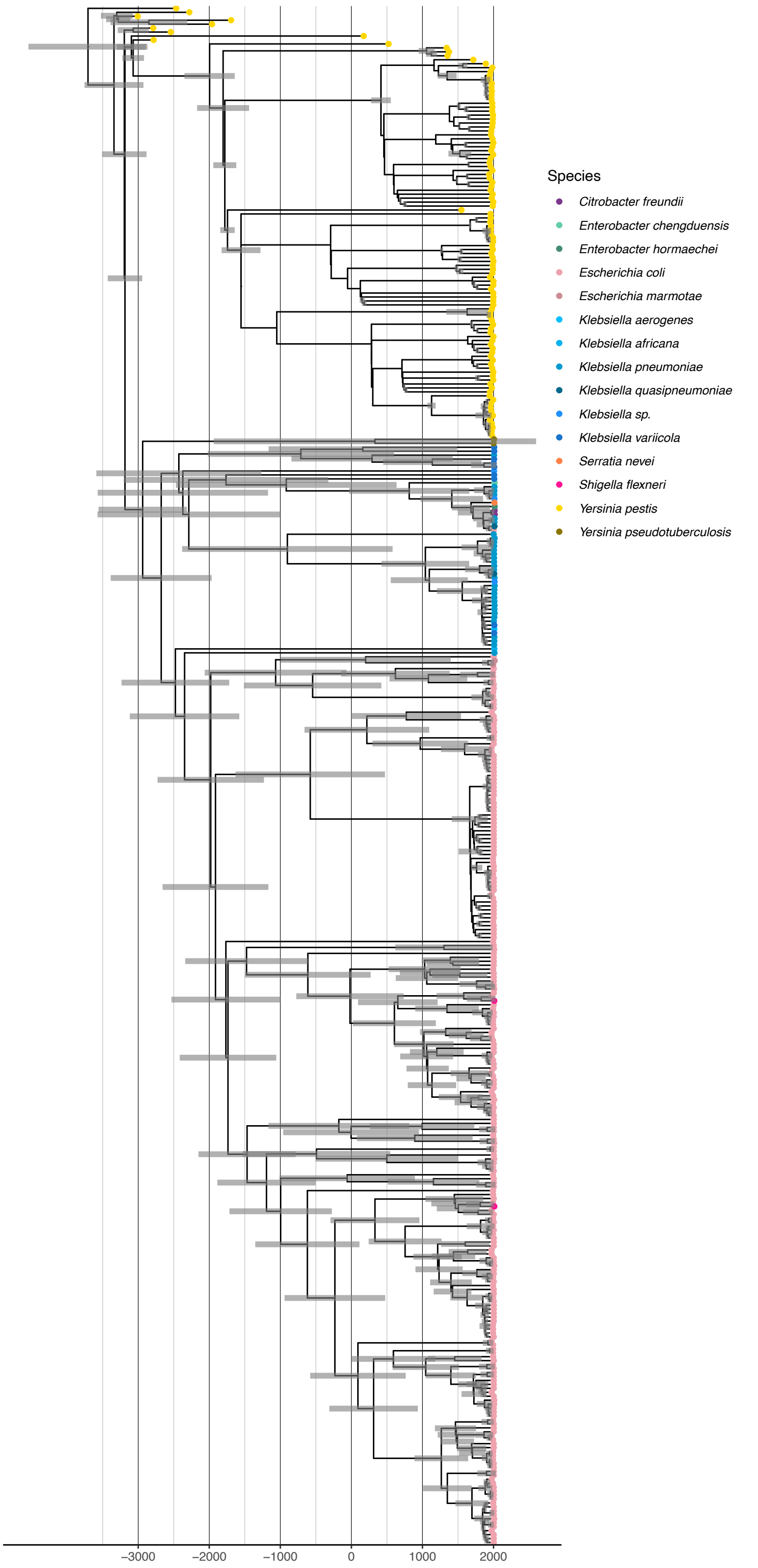


**Figure S5.** HPI dated tree rooted within *Y. pestis* (389 tips) with BEAST2, our favored scenario. HPD 95% confidence intervals for node ages are displayed as horizontal gray bars at each node.

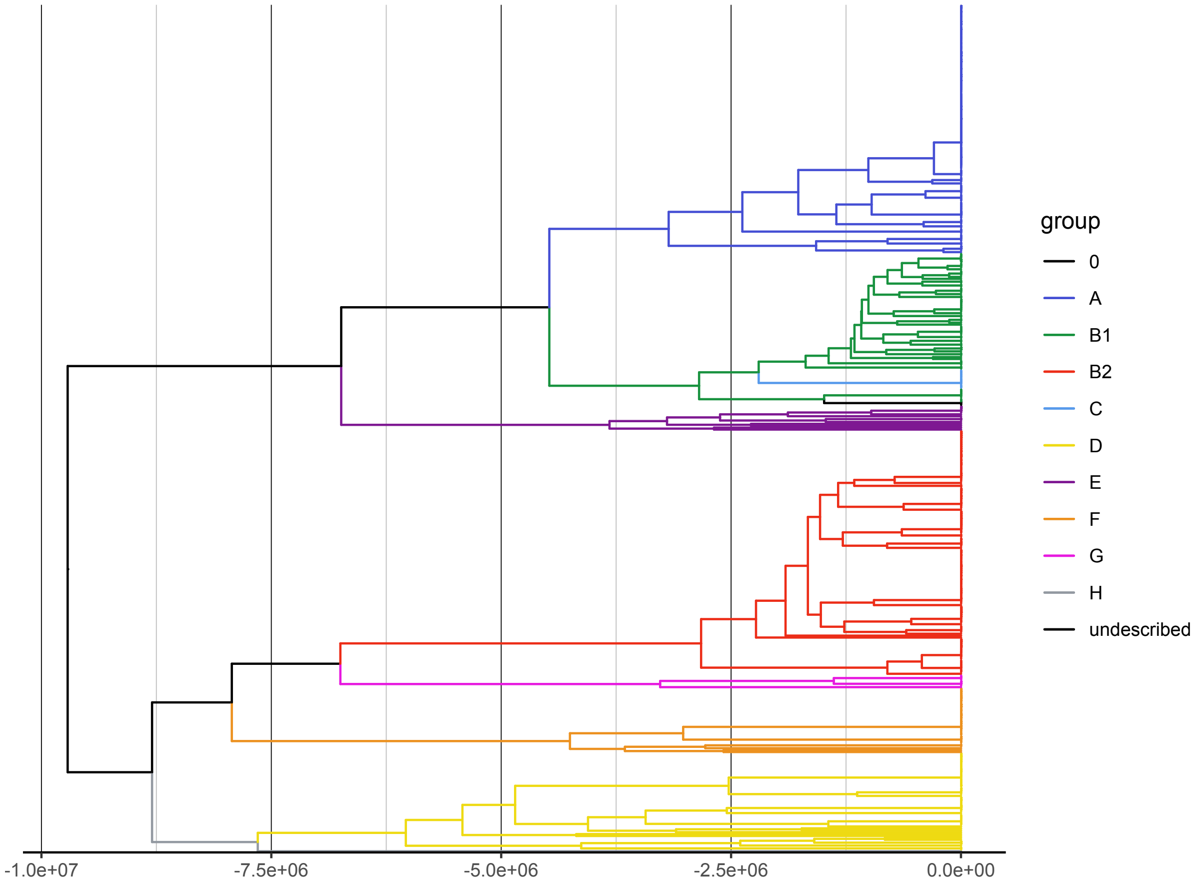


**Figure S6.** *E. coli sensu stricto* timetree (444 tips). Divergence times were inferred with the TDR method on the 450 strains. The three *S. enterica* and the three strains belonging to *E. ruysiae* (cryptic clade IV) and *E. marmotae* (cryptic clade V) were removed from this figure for more clarity.

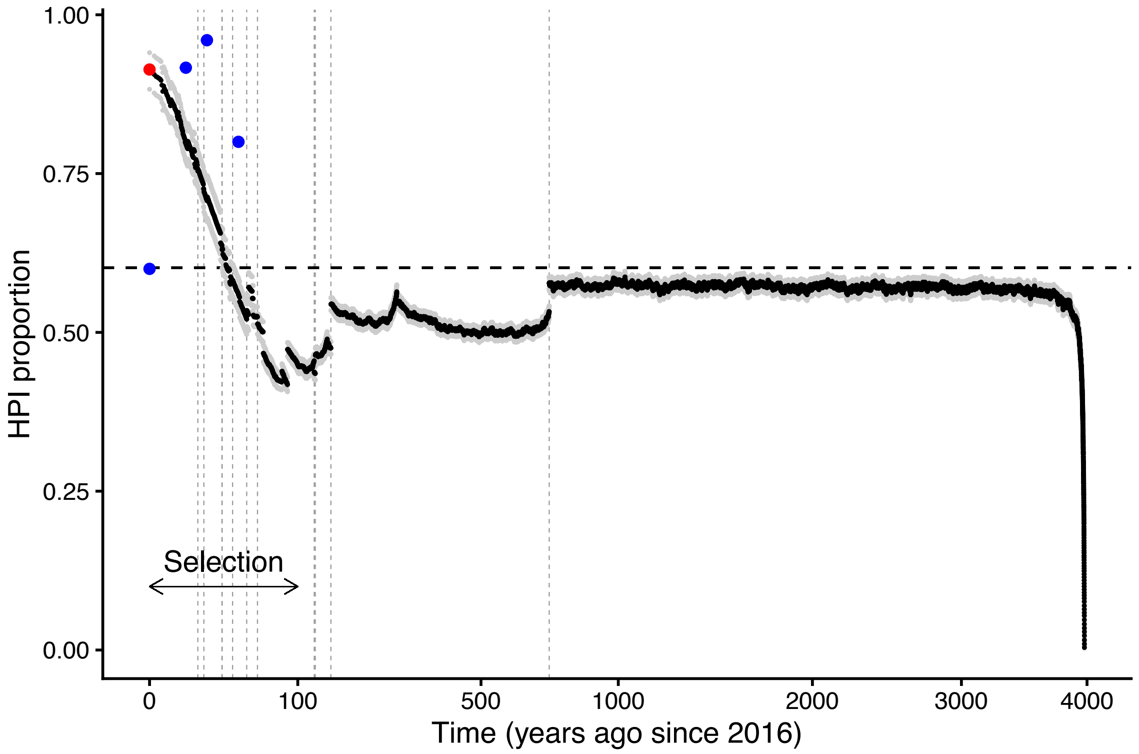


**Figure S7.** HPI proportion through time (with square root transformation of the x-axis) for B2 phylogroup. The presence of HPI along branches, since its introduction, was computed from 1000 character histories sampled from the posterior probability distribution (95%CI in gray). The dashed vertical gray lines indicate the time of the first branching event for each ancestral lineage. The dashed horizontal black line shows the expected ratio at equilibrium (q01/(q01+q10)). HPI frequency in B2 *E. coli* collections are illustrated by blues dots (CEREMI in 2016, Coliville in 2010, PAR/LBC/ROAR in 2001 and VDG in 1980). The HPI frequency at present time when considering all tips of the 11 B2 clades together is indicated by the red dot.

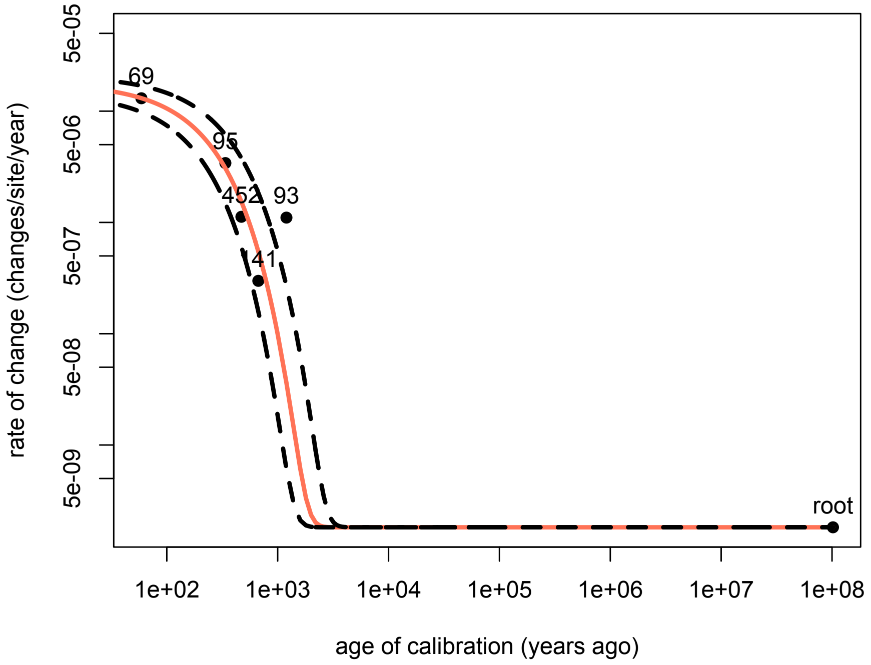


**Figure S8.** The rate of molecular change as a function of the age of calibration for the six STs with molecular clock signal (points). The point on the right at age 1e+08 years ago is the age of divergence between *E. coli* and *S. enterica.* We fitted a declining exponential curve to these six points (red line and 95% CI in dashed line). The black dashed lines represent the 95%CI. Both axes are in log scale.

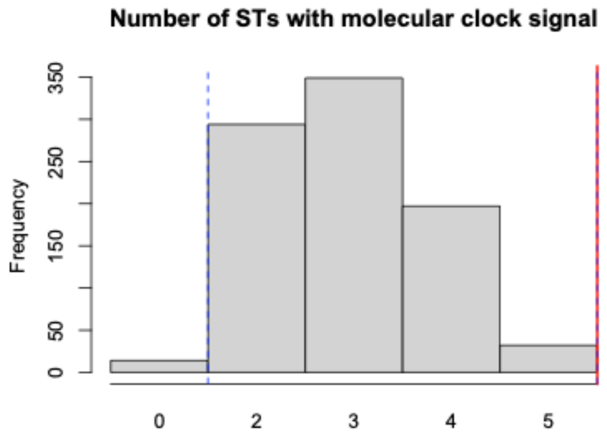


**Figure S9.** Null distribution of the number of STs with a molecular signal (*i.e.* positive relationship between root-to-tip distance and time). The null distribution was obtained by permutations of sampling dates among samples of a ST. The red line is the observed number of STs with a molecular clock signal. The dashed lines represent the 95% prediction interval.

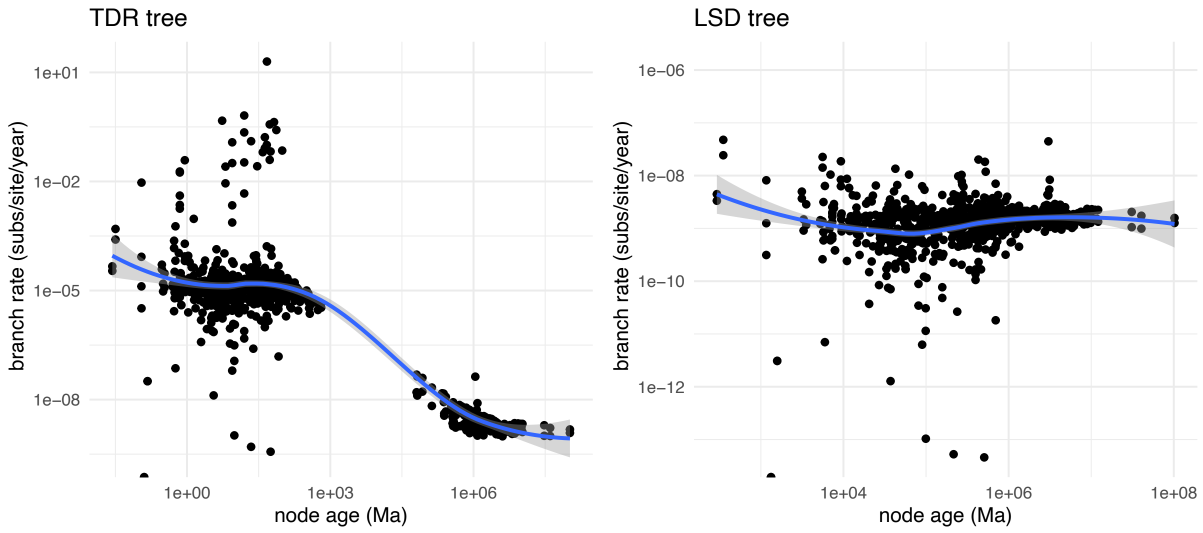


**Figure S10.** Molecular rate estimation (per branch, number of substitution/site/year) through time (ancestral node age, Ma) obtained with the TDR and LSD methods. Branch rates were obtained by dividing the branch lengths of the phylogenetic tree (fasttree tree) by the branch lengths of the timetrees (TDR and LSD trees). We fitted a Loess regression (blue line) with 95%CI (grey area).
